## Supplementary Figures 1-8 and Supplementary Table 1 for "Reciprocal impacts of telomerase activity and tumor cell differentiation in neuroblastoma tumor biology"

**Supplementary Table 1**

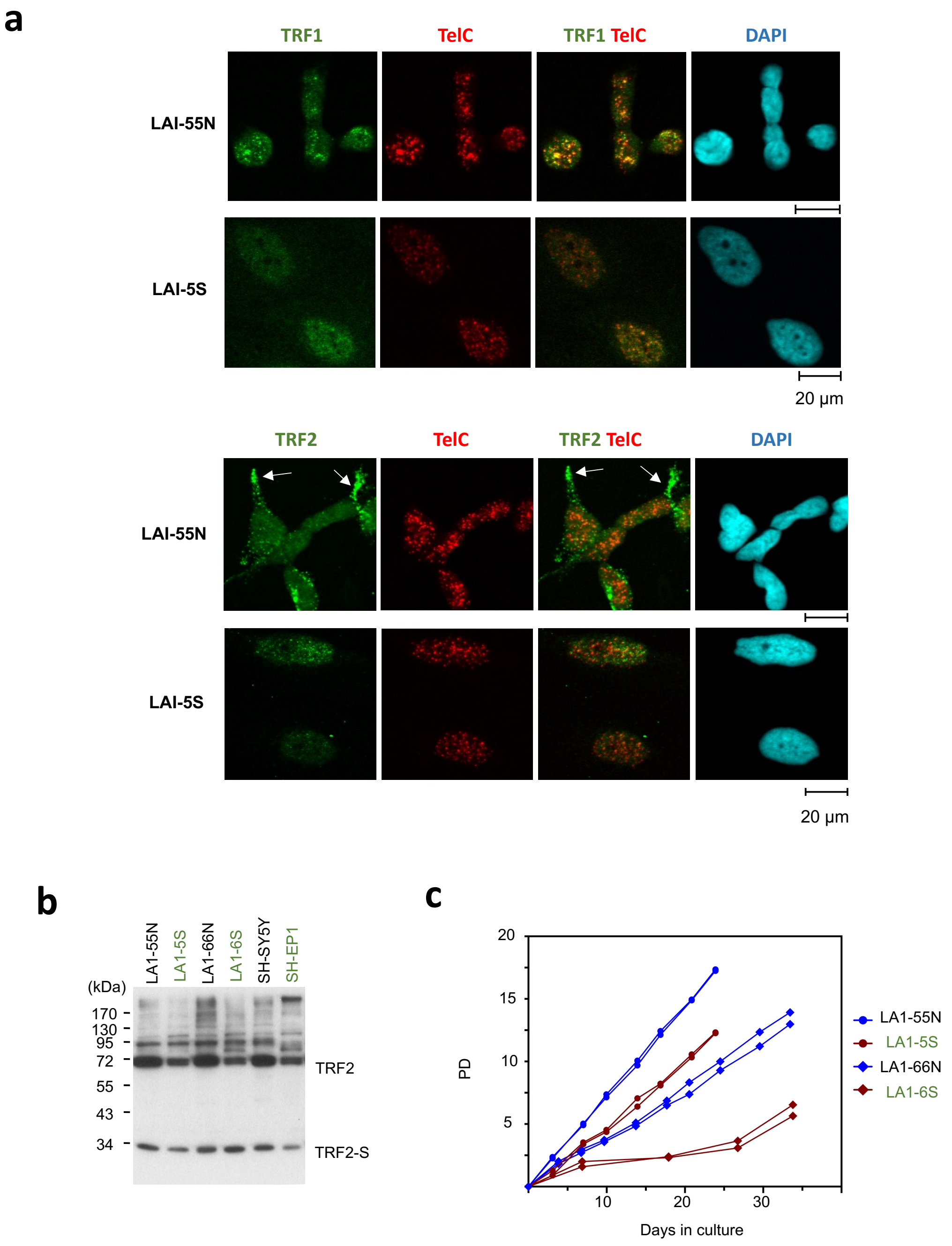


**Supp. Fig. 1. Differential cellular distribution of telomere protein in matched ADRN and MES cell lines**

(a) and (b) IF-FISH analysis of TRF1 and TRF2 in the indicated cell lines. Prominent examples of peri-nuclear staining of TRF2 in LA1-55N are marked white arrows.

(b) Western analysis of TRF2 and TRF2-S. TRF2-S  was detected by longer exposures of TRF2 Western blots.

(c) Comparison of growth rate in two matched ADRN and MES cell lines. The LA1-55N (ADRN) and LA1-5S (MES) pair and LA1-66N (ADRN) and LA1-6S (MES) pair were used.

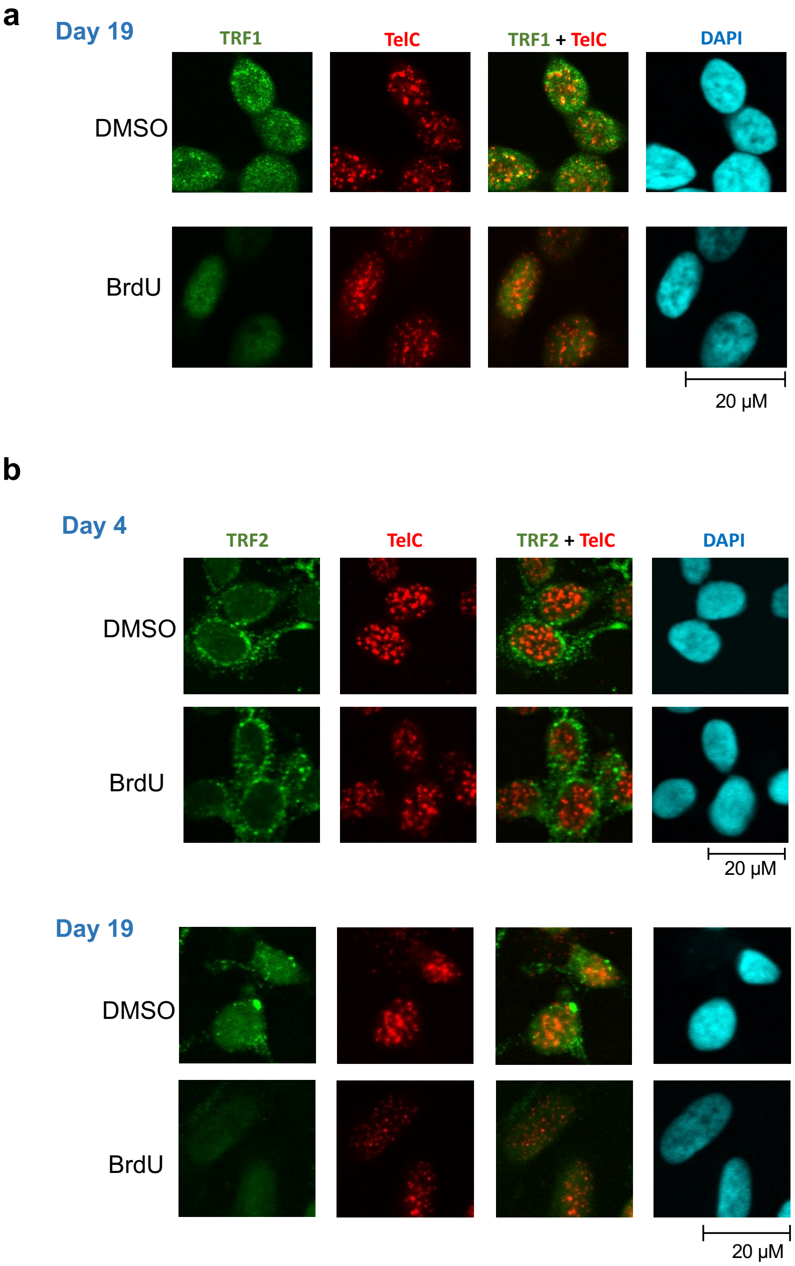


**Supp. Fig. 2. Cellular distribution of TRF1 and TRF2 in BE(2)N cells during BrdU-induced switch from ADRN to MES cell types**
(a) and (b) IF-FISH analysis of TRF1 and TRF2 in BE(2)N at different time points during BrdU-induced phenotypic switch.

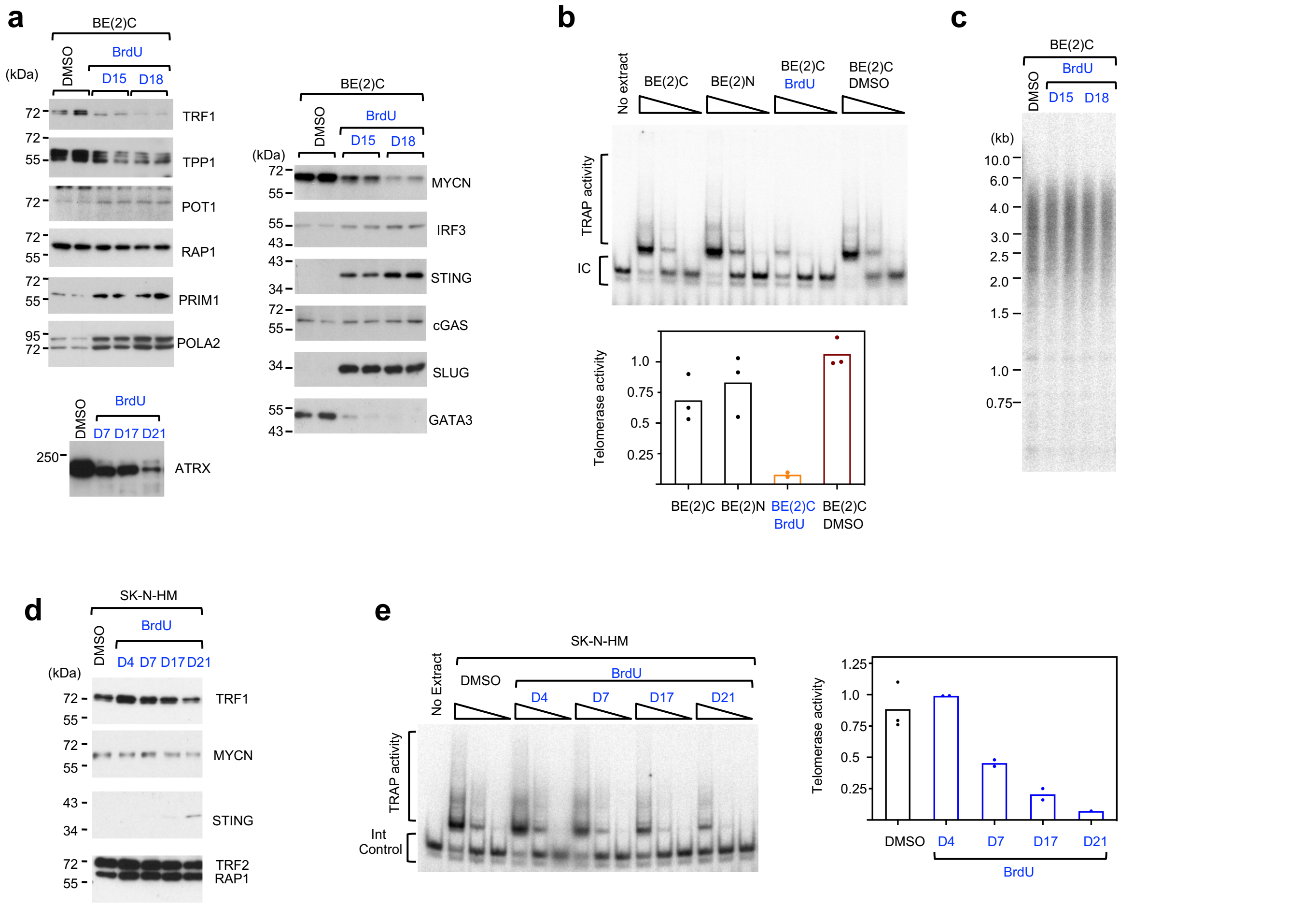


**Supp. Fig. 3. Profiling of BE(2)C and SK-N-HM cells during BrdU-induced switch from ADRN to MES cells**

(a) Western analysis of telomere and lineage-related and DNA sensing factors in BE(2)C during BrdU-induced phenotypic switch.

(b) TRAP analysis of telomerase activity in BE(2)C at different time points during BrdU-induced phenotypic switch. The assays were performed using serial dilutions of the extracts (equivalent to 500, 50 and 5 ng of proteins), and the relative activities are plotted at the bottom.

(c) Analysis of telomere length distributions in BE(2)C at different time points during BrdU-induced phenotypic switch.

(d) Western analysis of telomere and lineage-related and DNA sensing factors in SK-N-HM during BrdU-induced phenotypic switch.

(e) TRAP analysis of telomerase activity in SK-N-HM at different time points during BrdU-induced phenotypic switch. The assays were performed using serial dilutions of the extracts (equivalent to 500, 50 and 5 ng of proteins), and the relative activities are plotted.

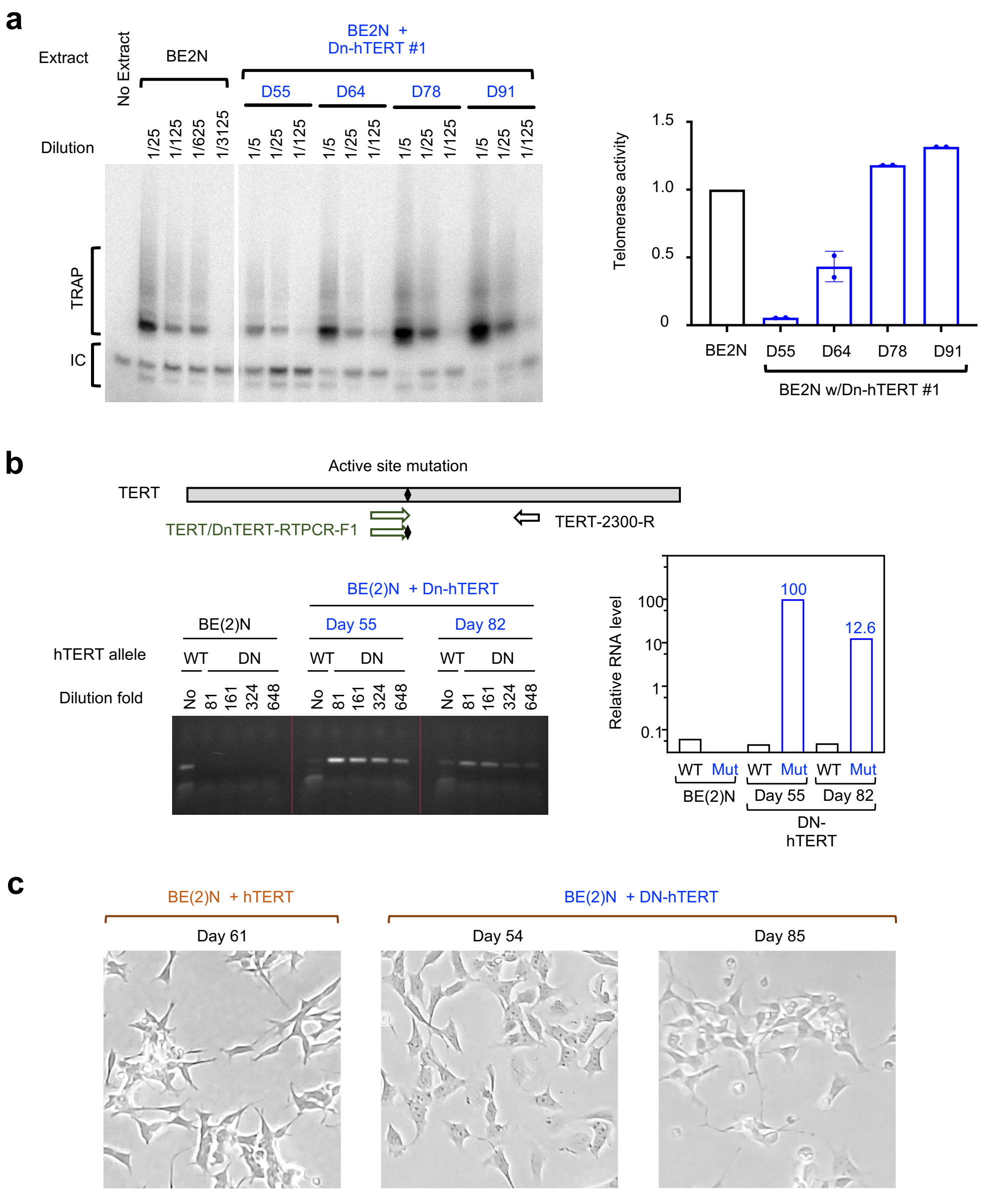


**Supp. Fig. 4. Characterization of BE(2)N cells during Dn-hTERT-induced switch from ADRN to MES cells**

(a) TRAP analysis of telomerase activity in BE(2)N harboring either hTERT or Dn-hTERT at different time points following retrovirus infection. The assays were performed using serial dilutions of the extracts as indicated. The relative activity of each extract was quantified using ImageQuant and plotted.

(b) Allele-specific RT-PCR analysis with total RNA from parental BE(2)N and BE(2)N harboring Dn-hTERT at different time points following retrovirus infection. Total RNAs were serially diluted as indicated and applied to the assays. The relative levels of wild type hTERT and DN-hTERT mRNA were quantified using ImageQuant and plotted.

(c) Morphology of BE(2)N harboring either hTERT or Dn-hTERT at different time points following retrovirus infection.

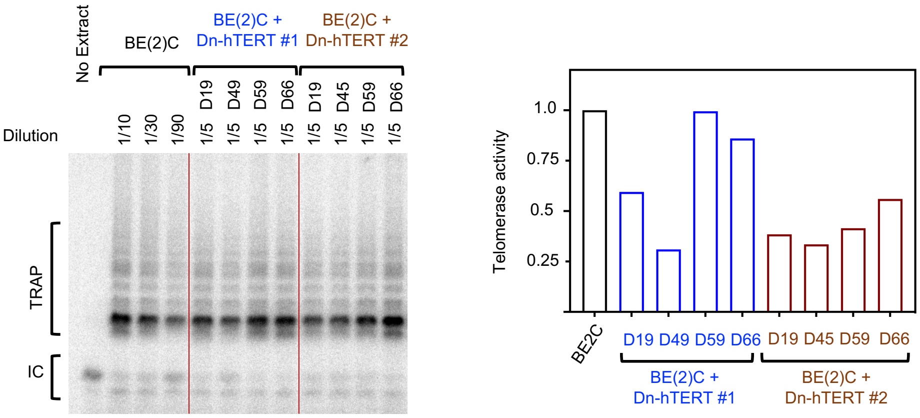


**Supp. Fig. 5. Mild inhibition of telomerase activity in Dn-hTERT treated BE(2)C**

TRAP analysis of telomerase activity in BE(2)C harboring Dn-hTERT at different time points following retrovirus infection. The assays were performed using different dilutions of the extracts as indicated. The relative activity of each extract was quantified using ImageQuant and plotted.

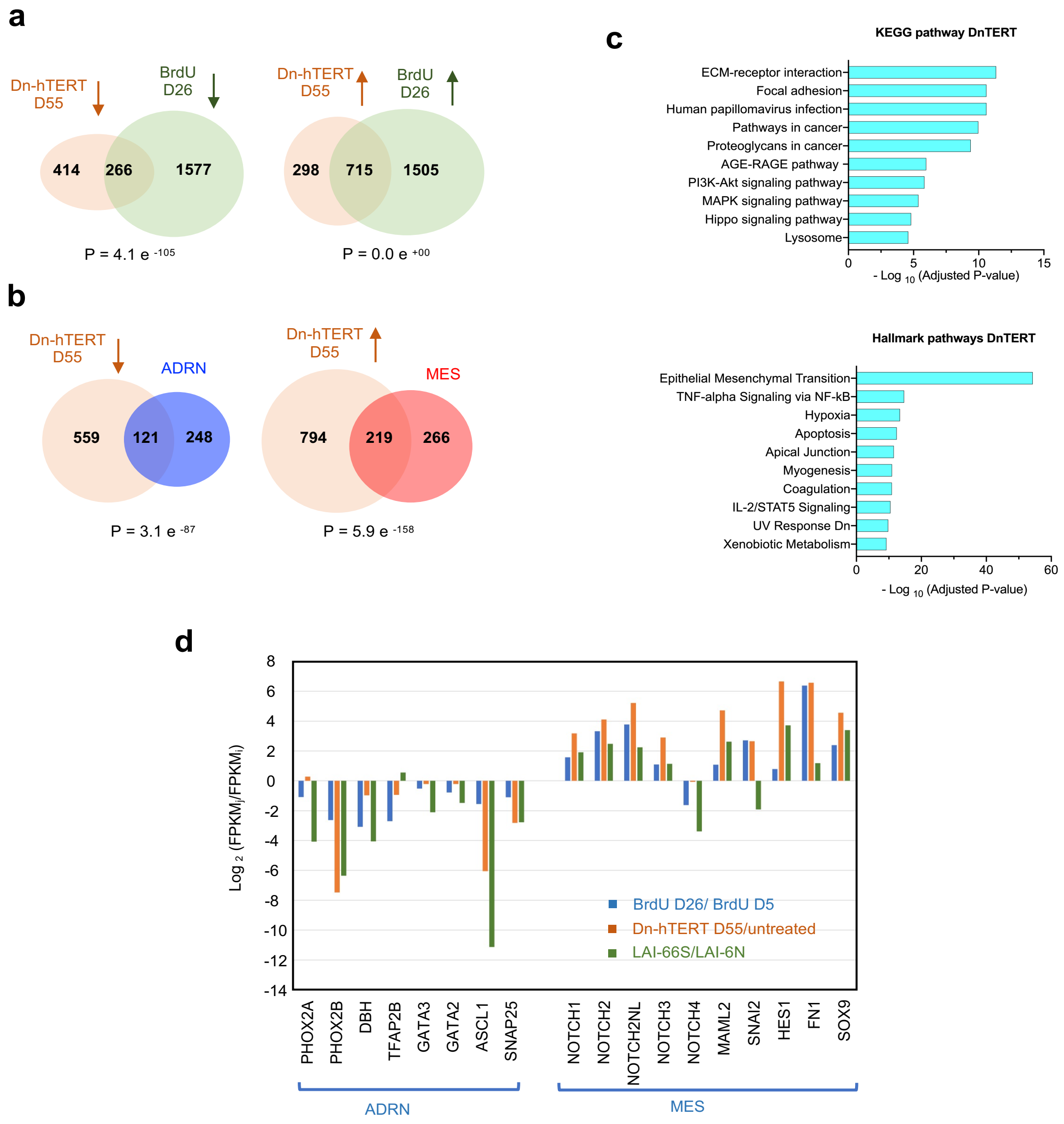


**Supp. Fig. 6. Comparison of gene expression pattern among Dn-hTERT-induced MES-like BE(2)N cells and BrdU-induced MES-like BE(2)N cells, and with previously defined MES and ADRN signature genes**

(a) Pair-wise comparison of the degree of overlaps in down-regulated genes (left) and up-regulated genes (right) between Dn-hTERT-induced MES-like BE(2)N cells (day 55) and BrdU-induced MES cells (day 26).

(b) (left) The list of genes that were down-regulated in Dn-hTERT-induced MES-like BE((2)N cells (day 55) (by >2-fold) was compared to the ADRN signature list [10]. (right) In parallel, the list of genes that were up-regulated in Dn-hTERT-induced MES-like BE(2)N cells (day 55) (by >2-fold) was compared to the MES signature.

(c) The ~1,000 genes up-regulated in Dn-hTERT-treated BE(2)N (by >2-fold) were subjected to pathway analysis by the Enrichr program. The top ten GO terms identified by the KEGG pathway database and the Hallmark gene sets in the Molecular Signature Database were ranked by adjusted P-values and plotted in the top and bottom panels, respectively.

(d) The expression changes of a subset of ADRN and MES signature genes in the BrdU experiment (day 26 vs day 5) and Dn-hTERT experiment (day 55 vs parental cell) are plotted. Also included are the relative expression of these genes between a matched pair of ADRN and MES cells (LA1-66N and LA1-6S).

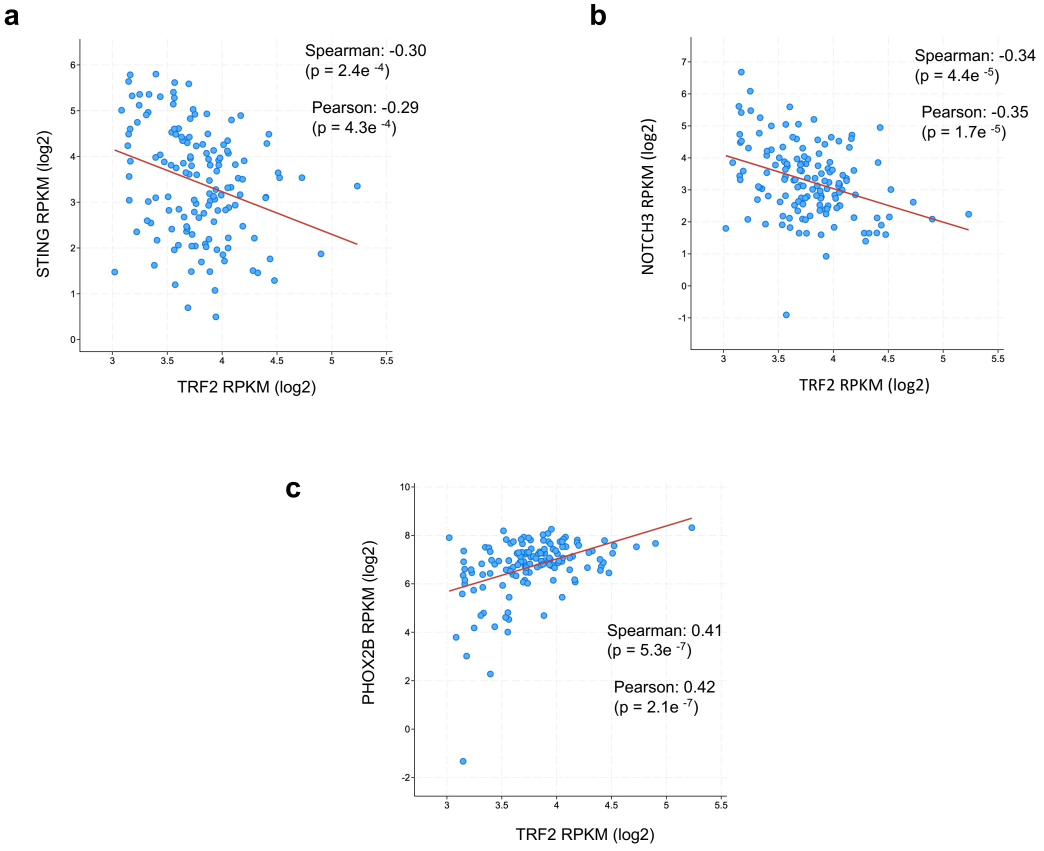


**Supp. Fig. 7. Correlations between the levels of TRF2 RNA and three other genes implicated in NB cell lineage regulation in patient samples**

The RNA levels of TRF2 in relation to three genes implicated NB cell lineage determination (i.e., STING [immunity], NOTCH3 [MES marker] and PHOX2B [ADRN marker]) in NB tumors from the Pediatric Neuroblastoma (TARGET, 2018) collection were analyzed and visualized using cBioportal.

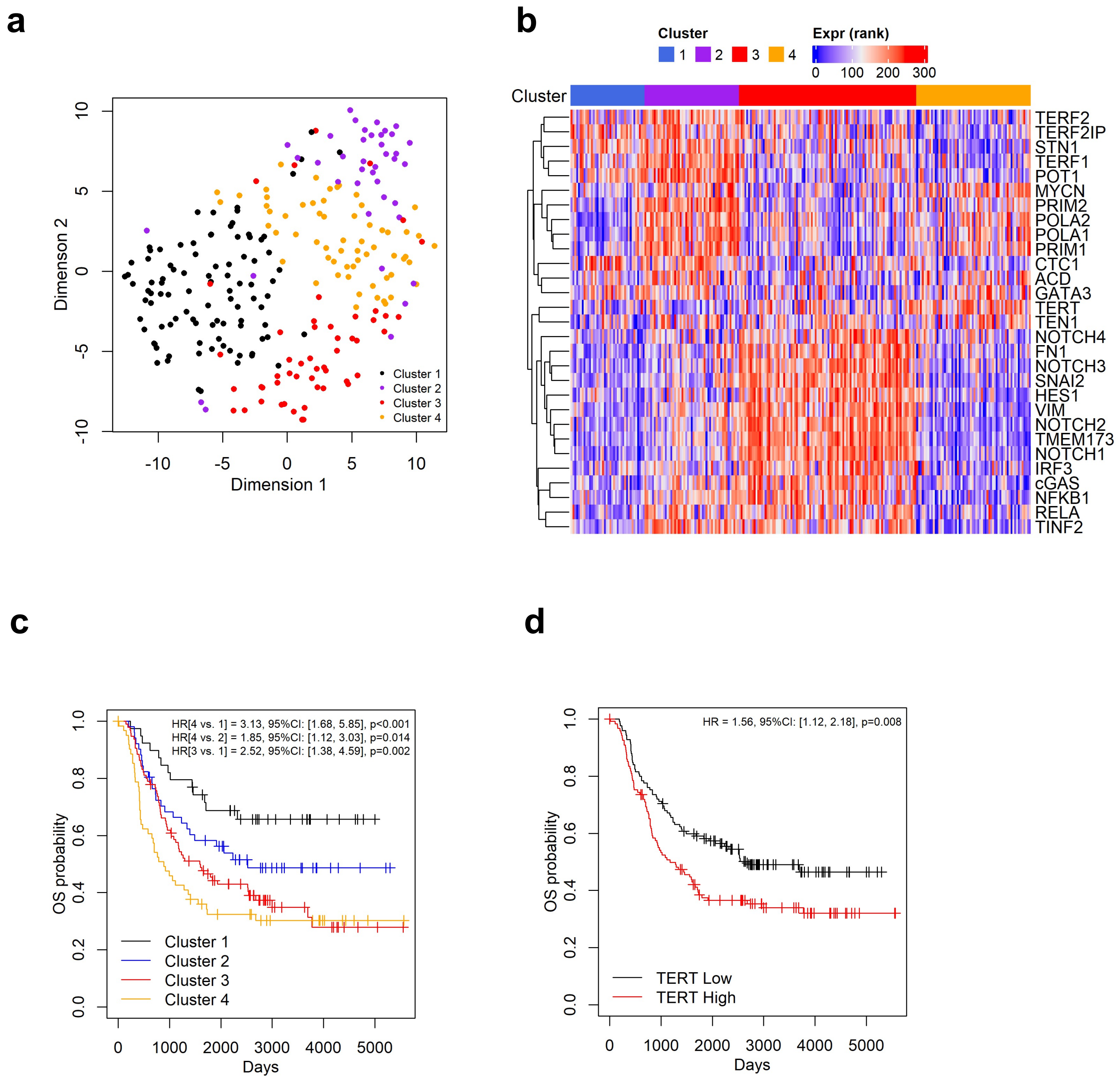


**Supp. Fig. 8. Analysis of the expression patterns and clinical outcomes associated with NB tumor samples with microarray data using a telomere- and cell lineage-related signature gene list**

(a) The RNA levels (rank transformed) of a list of 29 telomere-, cell lineage- and immunity-related genes across 247 NB tumors with microarray data were analyzed and clustered using PAM and displayed as a t-SNE plot.

(b) The rank transformed RNA levels of signature genes across four clusters of NB tumor samples are displayed in Heatmap.

(c) Kaplan-Meier overall survival curves for the four clusters of patients with distinct gene expression profiles.

(d) Kaplan-Meier overall survival curves for patients in the microarray dataset with high TERT and low TERT expression.

**Supp. Table 1. Gene expression differences between cluster 1 and cluster 3 patients**

| | All(n=85) | 1(n=35) | 3(n=50) | P-value* |
|---|---|---|---|---|
| **AGE** | | | | |
| Mean+/-sd | 3.92+/-2.52 | 2.91 +/- 2.29 | 4.62 +/- 2.46 | 0.002 |
| Median (IQR) | 3 (2, 5) | 2 (1,4) | 4 (3,6) | <0.001 |
| **INSS_STAGE, n (%)** | | | | |
| Stage 2b | 1(1.18%) | 1 (2.86%) | 0 (0%) | |
| Stage 3 | 4(4.71%) | 3 (8.57%) | 1 (2%) | |
| Stage 4 | 68(80%) | 20 (57.14%) | 48 (96%) | |
| Stage 4s | 12(14.12%) | 11 (31.43%) | 1 (2%) | <0.001 |
| **TUMOR_SAMPLE_HISTOLOGY, n (%)** | | | | |
| Favorable | 15(17.65%) | 12 (34.29%) | 3 (6%) | |
| Unfavorable | 66(77.65%) | 21 (60%) | 45 (90%) | |
| Unknown | 4(4.71%) | 2 (5.71%) | 2 (4%) | 0.001 |
| **RISK_GROUP, n (%)** | | | | |
| High Risk | 69(81.18%) | 20 (57.14%) | 49 (98%) | |
| Intermediate Risk | 9(10.59%) | 8 (22.86%) | 1 (2%) | |
| Low Risk | 7(8.24%) | 7 (20%) | 0 (0%) | <0.001 |
| **OS_STATUS, n (%)** | | | | |
| 0:LIVING | 37(43.53%) | 21 (60%) | 16 (32%) | |
| 1:DECEASED | 48(56.47%) | 14 (40%) | 34 (68%) | 0.015 |
| **OS_DAYS** | | | | |
| Mean+/-sd | 1557.44+/-1043.28 | 1826.94 +/- 1162.02 | 1368.78 +/- 916.92 | 0.056 |
| Median (IQR) | 1549 (659, 2325) | 2064 (806,2500) | 1323.5 (487.25,2012.75) | 0.04 |
| **OS_MONTHS** | | | | |
| Mean+/-sd | 51.68+/-34.28 | 60.46 +/- 38.14 | 45.54 +/- 30.19 | 0.058 |
| Median (IQR) | 51 (22, 77) | 68 (27,82.5) | 44 (16.75,67) | 0.042 |
| **TERT** | | | | |
| Mean+/-sd | 0.71+/-0.94 | 0.36 +/- 0.59 | 0.95 +/- 1.05 | 0.002 |
| Median (IQR) | 0.08 (0, 1.25) | 0 (0,0.78) | 0.65 (0.03,1.77) | <0.001 |
| **TERF2** | | | | |
| Mean+/-sd | 4.01+/-0.34 | 3.9 +/- 0.3 | 4.1 +/- 0.35 | 0.005 |
| Median (IQR) | 3.98 (3.81, 4.15) | 3.91 (3.75,4.04) | 4.05 (3.86,4.27) | 0.006 |
| **TERF2IP** | | | | |
| Mean+/-sd | 6.59+/-0.56 | 6.78 +/- 0.46 | 6.46 +/- 0.59 | 0.007 |
| Median (IQR) | 6.65 (6.32, 6.94) | 6.85 (6.52,7.04) | 6.55 (6.05,6.72) | 0.004 |
| **POLA1** | | | | |
| Mean+/-sd | 2.55+/-0.56 | 2.19 +/- 0.52 | 2.79 +/- 0.43 | <0.001 |
| Median (IQR) | 2.58 (2.25, 2.91) | 2.26 (1.92,2.5) | 2.73 (2.54,3.04) | <0.001 |
| **POLA2** | | | | |
| Mean+/-sd | 3.94+/-0.92 | 3.45 +/- 1.02 | 4.29 +/- 0.67 | <0.001 |
| Median (IQR) | 4.06 (3.5, 4.59) | 3.53 (2.69,4.21) | 4.23 (3.91,4.73) | <0.001 |

| | | | | |
|---|---|---|---|---|
| **STN1** | | | | |
| Mean+/-sd | 2.93+/-0.44 | 3.15 +/- 0.5 | 2.77 +/- 0.32 | <0.001 |
| Median (IQR) | 2.89 (2.65, 3.22) | 3.23 (2.9,3.42) | 2.79 (2.65,2.96) | <0.001 |
| **TEN1** | | | | |
| Mean+/-sd | 3.11+/-0.51 | 3.34 +/- 0.47 | 2.94 +/- 0.48 | <0.001 |
| Median (IQR) | 3.14 (2.72, 3.4) | 3.35 (3.14,3.55) | 2.95 (2.65,3.26) | <0.001 |

* P-values were determined using the ANOVA and the non-parametric Kruskal-Wallis tests.
